# Transcriptional mapping of the macaque retina and RPE-choroid reveals conserved inter-tissue transcription drivers and signaling pathways

**DOI:** 10.1101/2022.01.21.477280

**Authors:** Ameera Mungale, David M. McGaughey, Congxiao Zhang, Sairah Yousaf, James Liu, Brian P. Brooks, Arvydas Maminishkis, Temesgen D. Fufa, Robert B. Hufnagel

## Abstract

**Purpose:** The macula and fovea comprise a highly sensitive visual detection tissue that is susceptible to common disease processes like age-related macular degeneration (AMD). Our understanding of the molecular determinants of high acuity vision remains unclear, as few model organisms possess a human-like fovea. We explore transcription factor networks and receptor-ligand interactions to elucidate tissue interactions in the macula and peripheral retina and concomitant changes in the underlying retinal pigment epithelium (RPE)/choroid.

**Methods:** Poly-A selected, 100 bp paired-end RNA-sequencing (RNA-seq) was performed across the macular/foveal, perimacular, and temporal peripheral regions of the neural retina and RPE/choroid tissues of four adult Rhesus macaque eyes to characterize region- and tissue-specific gene expression. RNA-seq reads were mapped to both the macaque and human genomes for maximum alignment and analyzed for differential expression and Gene Ontology (GO) enrichment.

**Results:** Comparison of the neural retina and RPE/choroid tissues indicated distinct, contiguously changing gene expression profiles from fovea through perimacula to periphery. Top GO enrichment of differentially expressed genes in the RPE/choroid included cell junction organization and epithelial cell development. Expression of transcriptional regulators and various disease-associated genes show distinct location-specific preference and retina-RPE/choroid tissue-tissue interactions.

**Conclusions:** Regional gene expression changes in the macaque retina and RPE/choroid is greater than that found in previously published transcriptome analysis of the human retina and RPE/choroid. Further, conservation of human macula-specific transcription factor profiles and gene expression in macaque tissues suggest a conservation of programs required for retina and RPE/choroid function and disease susceptibility.

## Introduction

The retina is a thin tissue of the posterior eye (range: 166.9 ± 20.9 μm - 271.4 ± 19.6 μm[1] essential for processing light and transmitting information to the brain through the optic nerve. It is composed of multiple layers, which contain seven cell classes including rod and cone photoreceptors, in apposition with the retinal pigment epithelium (RPE), which is underlaid by the choroid. The choroid is a vascular structure that supports the outer retina, the RPE is necessary for maintaining healthy photoreceptors, and rods mediate dark-adapted vision while cones are responsible for color vision[2, 3]. The relative composition of cell types varies across the retina, and provides visual function specializations, such as high acuity color vision at the fovea, which is cone photoreceptor rich and rod depleted. Furthermore, the neural retinal layers change in thickness and morphology, including the inner and outer nuclear layers of the photoreceptors, whereas the choroidal layer has increased thickness at the macula but the morphology of the entire layer remains relatively unchanged[4, 5]. Many genes have varied expression in different cell types and tissue layers, with 819 genes implicated in heritable retinal degenerations as of 2022 (RetNet, https://sph.uth.edu/RetNet/)[6, 7]. While retinal degeneration may initiate in a single tissue layer, eventually the disease progresses across all tissue layers [8]. Importantly, retinal manifestations in many diseases are region specific. For example, Stargardt disease and Best disease primarily affect the macula and cause loss of central vision first, whereas retinitis pigmentosa initiates in the rod-rich periphery[9–12]. We hypothesize that this location-specific disease susceptibility of the retina is due to cellular specialization and region-dependent molecular interactions, which we can assess by regional layer-specific gene expression profiling of healthy retinae.

Previous investigations of the transcriptomic landscape of the human retina support a significant differential gene expression between the retina and the RPE/choroid/sclera tissue layers as well as across macular and peripheral regions, as described by Li et al., 2014[13]. Comparison of gene expression between the nasal and temporal peripheries of retina and RPE/choroid/sclera tissues indicated little to no variability, yet macular and peripheral profiles remained distinct in each tissue[13]. Gene expression across the retina has shown to be consistent with the spatial distribution of photoreceptors and ganglion cells, and RPE-specific genes appear to be enriched in the periphery over the macula as outlined by Whitmore et al., 2014[14]. Further characterization of gene expression between foveal and peripheral retina has been conducted using single-cell RNA sequencing in both primate (*Macaca fascicularis*) by Peng et al., 2019[15] and human retina by Voight et al., 2019[16]. These studies showed variability in both cell type distribution and gene expression between foveal and peripheral retina[15, 16]. While it has been well established that regional differences exist across tissue layers in the primate retina, the molecular drivers of these differences remain unclear.

Here we generate an RNA-seq dataset from adult Rhesus macaque (*Macaca mulatta*) eyes that reflect the changing photoreceptor composition of the contiguous macular/foveal, perimacular, and peripheral regions to determine tissue- and location-dependent differential gene expression of the neural retina and RPE/choroid. The use of matched retina and RPE/choroid tissue biopsies in this analysis with an additional perimacular sample provides a contiguous assessment of gene expression across location as opposed to a binary macular vs periphery comparison and allows for a more comprehensive look at patterns of gene expression and their related pathways and ontologies. We additionally explore conservation of these shared and distinct pathways through a meta-analysis comparing our macaque datasets to previously published human data. Finally, we examine gene regulation via transcription factors and tissue-tissue interactions using ligand/receptor analysis to understand differences in the macular and peripheral retina and the affiliated changes in the RPE/choroid.

## Methods

### Animal Care

All experimental protocols were approved by the National Eye Institute Animal Care and Use Committee. Procedures were performed in accordance with the United States Public Health Service policy on the humane care and use of laboratory animals.

### Tissue Acquisition

Postmortem eyes were enucleated from two adult Rhesus monkeys (*Macaca mulatta*), aged 13 and 18 years. Two-millimeter punch biopsies were removed from macular/foveal, perimacular, temporal, and nasal regions of the retina. Following the biopsy, the tissues were separated into layers of neural retina and RPE/choroid using microdissection techniques. Biopsies were optimized for smallest diameter in other macaque samples in order to obtain the minimal number of cells required for RNA-sequencing. Optimization was performed by measuring total RNA yield in different sized biopsy samples as below and requiring a minimum of 200 ng as input for RNA-seq.

### RNA Extraction

Total RNA was isolated from fresh tissues using the PicoPure™ RNA Isolation Kit (Applied Biosystems, Waltham, Massachusetts, USA) following mortar and pestle tissue homogenization. Extracted RNA samples were checked for quality using the 2100 Bioanalyzer (Agilent Technologies, Santa Clara, CA, USA) and only samples with RIN value > 6.5 were used (Range 1 – 10).

### RNA sequencing

Extracted RNA was prepared into 100 bp libraries for paired-end, poly-A-selected RNA sequencing on the Illumina HiSeq4000 at the NIH Intramural Sequencing Center. The raw sequence data is available at the Gene Expression Omnibus under accession GSE194285.

### Bioinformatic Analysis

Briefly, reads were quantified against the macaque genome Mmul_8.0.1 using Salmon pseudo-alignment transcript quantification tool (version 0.13.0; Patro et al., 2017)[17], imported into R with tximport[18] and analyzed for differential expression using DESeq2, using the Rhesus macaque sample as a covariate [19]. Transcript reads were additionally aligned to the human genome (GRCH38, gencode version 30 [20], to supplement any unquantified genes. To select the gene name for each macaque transcript, we used the HGNC Comparison of Orthology Predictions[21] data, aggregated by the hcop R package (https://github.com/stephenturner/hcop). We counted the number of sources supporting each gene ID to transcript and selected the gene ID with the most support, as detailed in https://github.com/davemcg/macaque_macula_RNA-seq/blob/master/analysis/01_QC_prelim_analysis.Rmd.

Genes were considered to be differentially expressed based on IHW[22] corrected p value < 0.05. Gene lists were then assigned gene ontologies using the clusterProfiler “enrichGO” function[23].

For the heatmap visualizations, we used length-scaled transcripts per million (TPM) quantification from tximport scaled by library size with the edgeR[24] “calcNormFactors” function. The heatmaps were made with the R package ComplexHeatmap[25].

Human single cell transcriptome data from the scEiaD resource at plae.nei.nih.gov was searched for machine predicted RPE cells from fovea or peripheral punches. We found 217 cells, 159 from the fovea and 58 from the periphery, across four studies[26–30]. The Seurat object was downloaded on 2022-01-14 from plae.nei.nih.gov and subsetted to these 217 cells. The Seurat object was converted to a SingleCellExperiment object to run wilcox and t tests with the scran “findMarkers” test and used the study as a covariate. Genes were kept which had both a wilcox AUC (area under curve) greater than 0.7 (1 is best), an abs(log2)fold change greater than 0.7, and a FDR corrected p value < 0.01. For the overlap testing we kept genes that had a padj < 0.01 in each comparison (single cell human RPE, bulk human RPE-choroid, or bulk macaque RPE-choroid) and the direction of the fold change between fovea and peripheral was the same. The venn diagram was created with the R package eulerr (https://cran.r-project.org/web/packages/eulerr/citation.html).

For the exact commands used in the analysis of this data, we make our Snakemake[31] reproducible workflow available at https://github.com/davemcg/macaque_macula_RNA-seq.

## Results

We performed RNA-seq on 2 mm retina and RPE/choroid tissues of 4 adult Rhesus macaque eyes in each of the central macular (foveal), perimacular, and peripheral regions (Fig. S1). Tissue dissections and RNA extractions were optimized for minimal input samples (<1 μg) to maximize spatial resolution. Principle component analysis (PCA) visualization indicates that samples group primarily by tissue. Foveal and peripheral samples generally cluster furthest from each other, and perimacular samples cluster in between (Fig. S2). We then assessed overall gene expression by tissue layer and location as well as differential gene expression between foveal samples and other peripheral samples (Table 1).

**Table 1.** A summary of the differential gene expression (padj < 0.05) in retina and RPE/choroid by location.

| Tissue | Location | # Genes |
| --- | --- | --- |
| Retina | Fovea | 6,921 |
|  | Perimacula | 6,926 |
|  | Periphery | 10,517 |
|  | Fovea vs Perimacula | 4,633 |
|  | Fovea vs Periphery | 9,644 |
| RPE/Choroid | Fovea | 6,775 |
|  | Perimacula | 7,465 |
|  | Periphery | 10,983 |
|  | Fovea vs Perimacula | 414 |
|  | Fovea vs Periphery | 7,925 |

Analysis of the foveal versus peripheral regions in the retina and RPE/choroid depicted distinct sets of differentially expressed genes in each tissue layer, with 9644 and 7925 differentially expressed genes in the retina and RPE/choroid, respectively (Padj < 0.05) (Table 1). Many of the significantly differentially expressed genes are known to have tissue-specific roles in development and disease (Fig. 1A, 1B). Retinal diseases also manifest uniquely at particular regions of the RPE or retina. For example, Age-Related Macular Degeneration (AMD) is thought to begin in the macular RPE[32], Late Onset Retinal Degeneration manifests in the peripheral RPE[33], Retinitis Pigmentosa causes photoreceptor loss in the peripheral retina[10], and finally, Cone-Rod Dystrophies primarily affect photoreceptors in the central retina[34].

**Figure 1.**
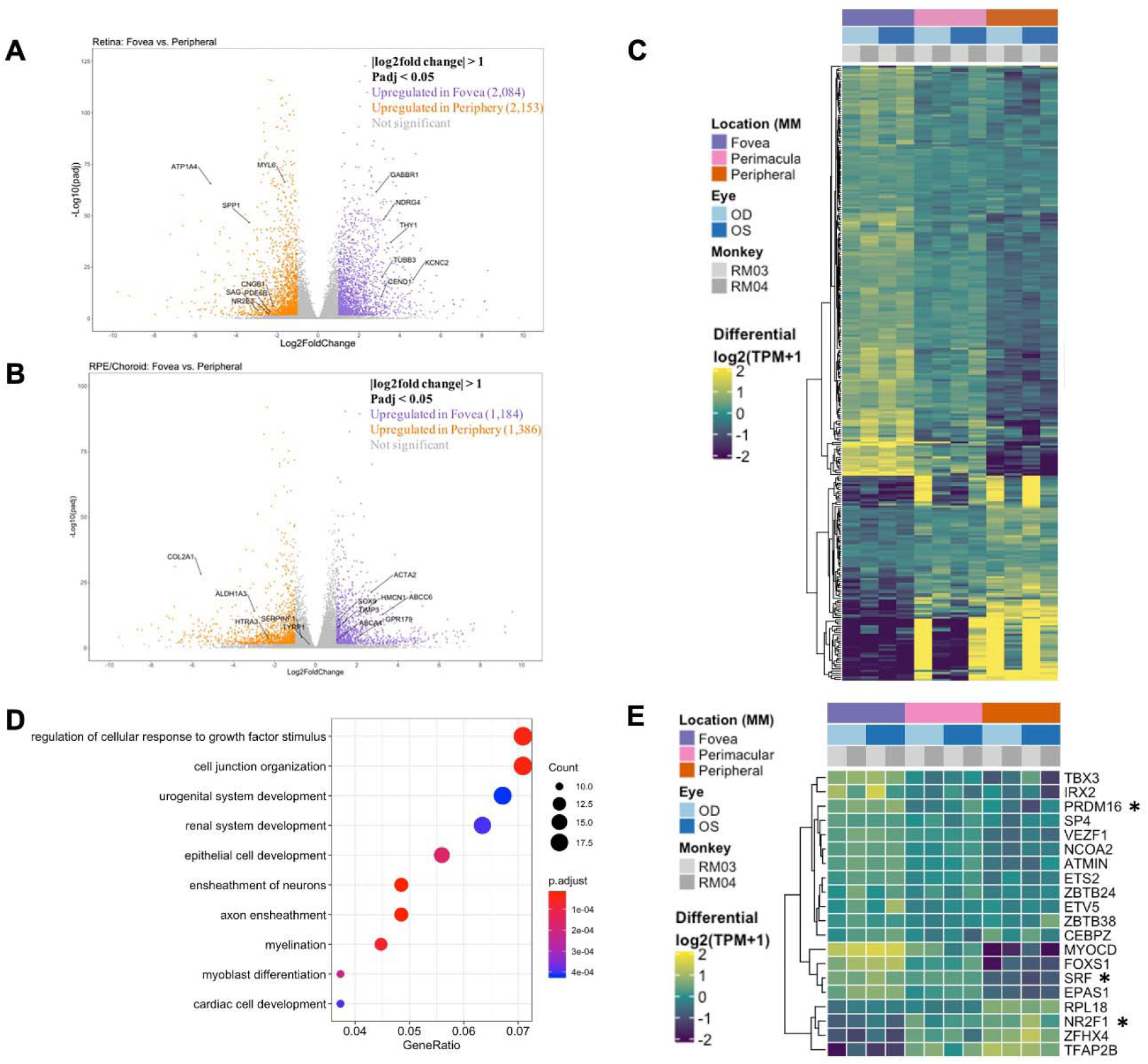
(A) A volcano plot depicting differentially expressed genes between foveal and peripheral tissues in the neural retina. Colored genes show significant up- or down-regulation with several labeled genes having known tissue-specific roles in development and disease including PDE6B. (B) A volcano plot showing RPE fovea vs periphery differential expression, with key genes labeled, including ABCA4 and TIMP3. (C) Unfiltered differential expression analysis of contiguously changing genes from fovea to perimacula to periphery in the RPE/choroid exhibits distinct blocks of lowly expressed genes in the fovea with a progressive increase in expression in peripheral tissues as well as blocks of genes with progressive downregulation moving outwards through the tissue despite less obvious morphologic changes in the tissue as compared to neural retina. (D) Top 10 enriched gene ontology (GO) terms from the RPE/choroid differentially expressed gene set include cell junction organization and epithelial cell development. VEGF is enriched in multiple of these ontologies and has increased expression in the macula over the periphery. (E) A subset of genes identified as differentially expressed transcription factors by location in the RPE/choroid. Changes in transcriptional regulators can be a factor in driving gene expression changes. *These transcription factors have published associations with the respective tissue in the literature.

In order to confirm that the differentially expressed genes across locations further reflect cellular composition of different tissues, we used expression patterns of published rod-enriched genes as identified in Holt et al., 2015[6] and Mustafi et al., 2016[35]. We observed correlated gene expression between eyes and animals, as well as a progressive increase in rod-enriched gene expression from the fovea to the perimacula and then to the periphery, reflecting changes in neural retina composition (Fig. S3A). In this list of rod enriched genes, we find that our peripheral retina samples are 2.8 fold enriched in rod signal relative to the fovea (t test p < 3.5 e-12). The perimacular region was enriched in rod signal by 1.7 fold over the fovea (p < 1.1 e-09). The enrichment of cone gene expression between the fovea and peripheral retina samples was 1.6 fold (p < 0.02) (Fig. S3B, Table S1). Among the RPE samples the enrichment of rod and cone markers was less substantial and not significant (Figure S3C).

Next, we performed an unfiltered analysis of the differentially expressed, contiguously changing genes from fovea to perimacula to periphery in the RPE/choroid. This analysis highlights stepwise changes due to changing photoreceptor composition in the cone-predominant macula, rod-predominant periphery, and admixed perimacula, and how these morphological changes in the neural retina may affect changing gene expression in the RPE/choroid. Differential expression analysis revealed distinct blocks of genes with low expression in the fovea that increase moving outward through the perimacula to the periphery, and other genes that are highly expressed in the fovea but progressively decrease moving outwards (Fig. 1C). We observe enrichment for gene ontologies including cell junction organization and epithelial cell development, as well as urogenital and renal system development (Fig. 1D).

Next, we sought to understand the transcriptional regulators driving the spatial and inter-tissue differences in gene expression. Using a list of all known transcription factors in humans, we identified a subset which is differentially expressed by location in each of the retina (Fig. S4B) and RPE/choroid datasets (Fig. 1E). After conducting a literature review, we further identified transcription factors with known associations in the respective tissues. In the retina and RPE, distinct sets of fovea-enriched and periphery-enriched transcription were elucidated. Our data shows three transcription factors, *NFRF1*, *IRX2*, and *ZFHX4*, to be differentially expressed in both neural retina and RPE/choroid and show a similar pattern in independent human data curated from eyeIntegration[36] (Fig. S5). Interestingly, *ZFHX4* appears to be enriched in the macular neural retina and the peripheral RPE/choroid, whereas *NFRF1* and *IRX2* expression is present in both tissues.

We then analyzed tissue-tissue interactions using ligand and receptor characterizations from the CellPhoneDB database[37]. After converting protein names from the database to gene names via the Uniprot conversion tool and merging the data with our location-based differential expression, we generated lists of distinct up- and down-regulated interactions in which both interacting partners in each tissue were either up- or downregulated in the fovea over the periphery (Table 2). Pathway analysis on these gene lists shows enrichment in the upregulated interactors for cell adhesion pathways (Fig. 2A, C) and that downregulated interactors were highly enriched for Wnt signaling in kidney disease and the ErbB signaling pathway, both of which are heavily involved in tissue development, including ocular development (Fig. 2B, D, Fig. S7C-D).

**Table 2.** Ligand-Receptor interactions across neural retina and RPE/choroid in the macula.

| Upregulated Interactions in the Macula |  | Downregulated Interactions in the Macula |  |
| --- | --- | --- | --- |
| RPE/Choroid | Neural Retina | RPE/Choroid | Neural Retina |
| NOTCH2 | JAG2 | EPHB2 | EFNB1 |
| FGFR1 | NCAM1 | WNT2B | FZD4 |
| NTF3 | NTRK3 | EFNA1 | EPHA2 |
| IGF2 | IGF1R | EPHB1 | EFNB1 |
| EPHA2 | EFNA3 | CCR1 | CCL26 |
| NOTCH1 | JAG2 | NRG2 | ERBB3 |
| PLXNB2 | SEMA4D | CD48 | CD244 |
| NTF3 | NTRK2 | FLT3 | FLT3LG |
| TNFSF10 | TNFRSF10D | DSC1 | DSG2 |
| EPHA3 | EFNA3 | EFNA1 | EPHA3 |
| TEK | ANGPT1 | PTPRC | CD22 |
| VEGFA | FLT1 | CCR5 | CCL8 |
| PLXNB2 | SEMA4G | LGALS9 | HAVCR2 |
| NOTCH3 | JAG2 | MDK | ALK |
| WNT1 | FZD1 | PLXNC1 | SEMA7A |
| NRP1 | VEGFA | ANXA1 | FPR2 |
| EGFR | EGF | FGFR1 | NCAM1 |
| EPHA4 | EFNB1 | CCR1 | CCL26 |
| NRP2 | SEMA3C | NGF | NTRK1 |
| ERBB4 | HBEGF | CD48 | CD244 |
| IGF2 | IGF2R | FZD7 | WNT3 |
| EFNA1 | EPHA2 | CCR6 | CCL20 |
| EPHB1 | EFNB1 | CCR2 | CCL26 |
| TNF | TNFRSF1A | CCR2 | CCL8 |
| IGF2 | IGF2R | PLXNB2 | SEMA4C |
| EFNA3 | EPHA4 | PVR | CD96 |
| CD55 | ADGRE5 | NRG1 | ERBB4 |
| CD44 | SELE | ERBB3 | NRG1 |
| ERBB4 | NRG4 | PTPRC | CD22 |
| CADM3 | CADM1 | SEMA5A | PLXNB3 |
| EFNA1 | EPHA3 | CSF1R | IL34 |
| CADM3 | EPB41L1 | CCR5 | CCL8 |
| VEGFA | FLT1 | PDCD1 | PDCD1LG2 |
| VEGFA | KDR | CCR1 | CCL8 |
| NRP2 | PGF | EREG | ERBB4 |
| DPP4 | CXCL12 |  |  |
| EPHB6 | EFNB1 |  |  |
| NRP2 | VEGFA |  |  |
| FLT1 | PGF |  |  |
| EFNA1 | EPHA4 |  |  |
| NRP2 | SEMA3F |  |  |

**Figure 2.**
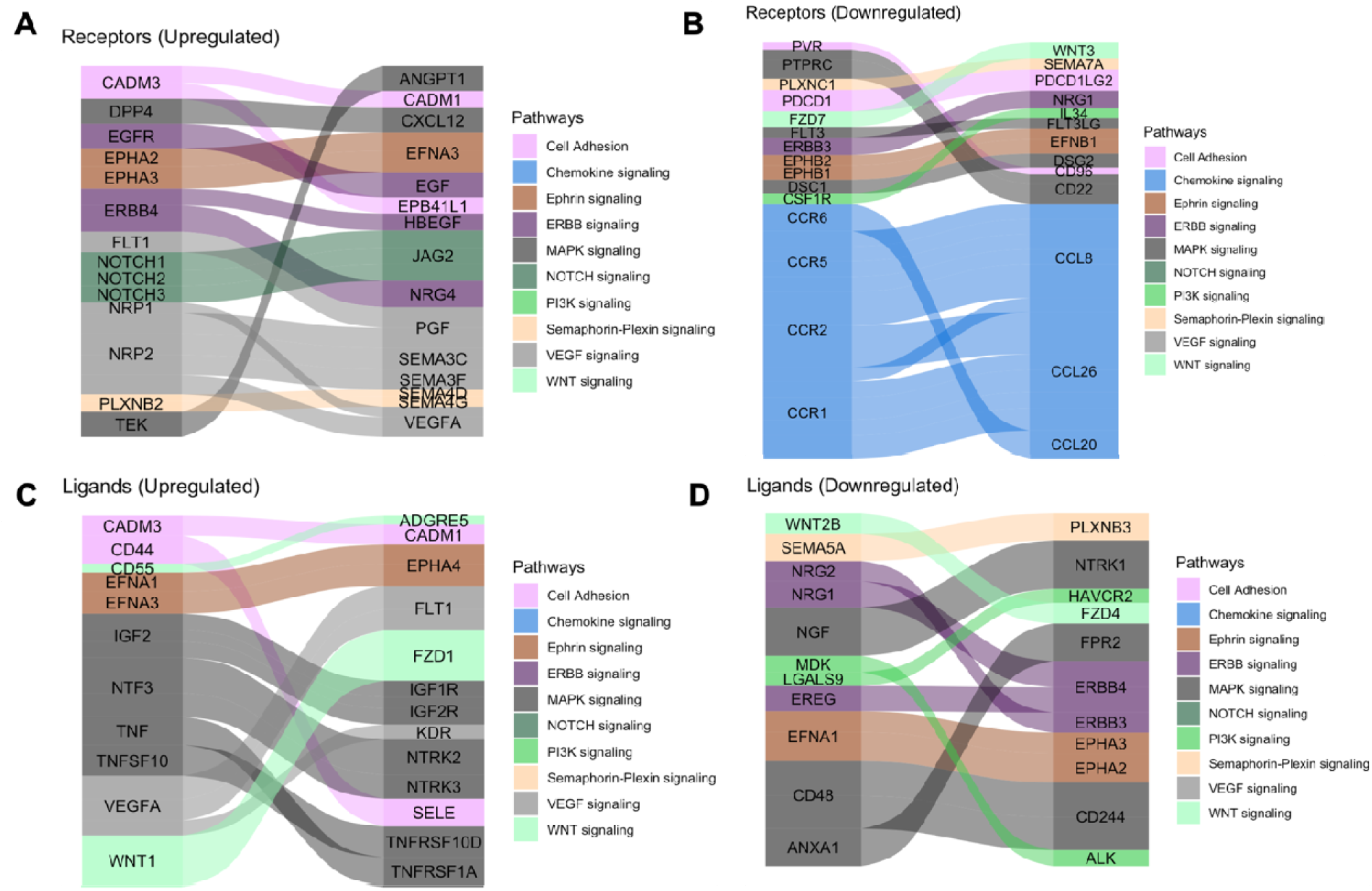
Tissue-tissue interactions showing receptor/ligand pairs in the central retina. (A) Upregulated macular receptors in the RPE/choroid are linked to their upregulated ligand in the macular neural retina. Interactors are grouped by signaling pathway and the width is scaled by average log2 fold change. (B) Downregulated macular receptors in the RPE/choroid and their respective downregulated ligand in the macular neural retina. (C) Upregulated macular ligands in the RPE/choroid are linked to their upregulated receptor in the macular neural retina. (D) Downregulated macular ligands in the RPE/choroid and their corresponding downregulated receptors in the macular neural retina.

To determine the conservation of location-specific differential expression of transcriptional regulators across fovea to periphery, we performed a meta-analysis comparing our macaque data to previously published human datasets[13, 14]. We see generally conserved patterns of expression from fovea to periphery in neural retina and to a lesser extent, RPE/choroid (Fig. 3A-B). We performed the same analysis on unfiltered gene lists from the human and macaque and found approximately 700 genes that follow similar differential expression patterns from fovea to periphery in both species. (Fig. 3A, Table 1S). GO terms for this list of conserved genes involve neuronal connectivity and cellular morphogenesis (Fig. 3C), similar to the GO terms for the macaque retina data seen previously in this study. In one case, we see a distinct block of macula-specific transcription factors in the Rhesus monkey neural retina including *HR*, *MYC*, *EBF3*, *AR*, *EBF2*, *EBF1*, *TWIST1*, *IRX2*, *POU4F2*, *IRX1*, and *SHOX2* that seem to also correlate to macula-specific transcription factors in the human data (Fig. S6B). Overall, this supports a conservation of gene expression and function across monkey and human species.

**Figure 3.**
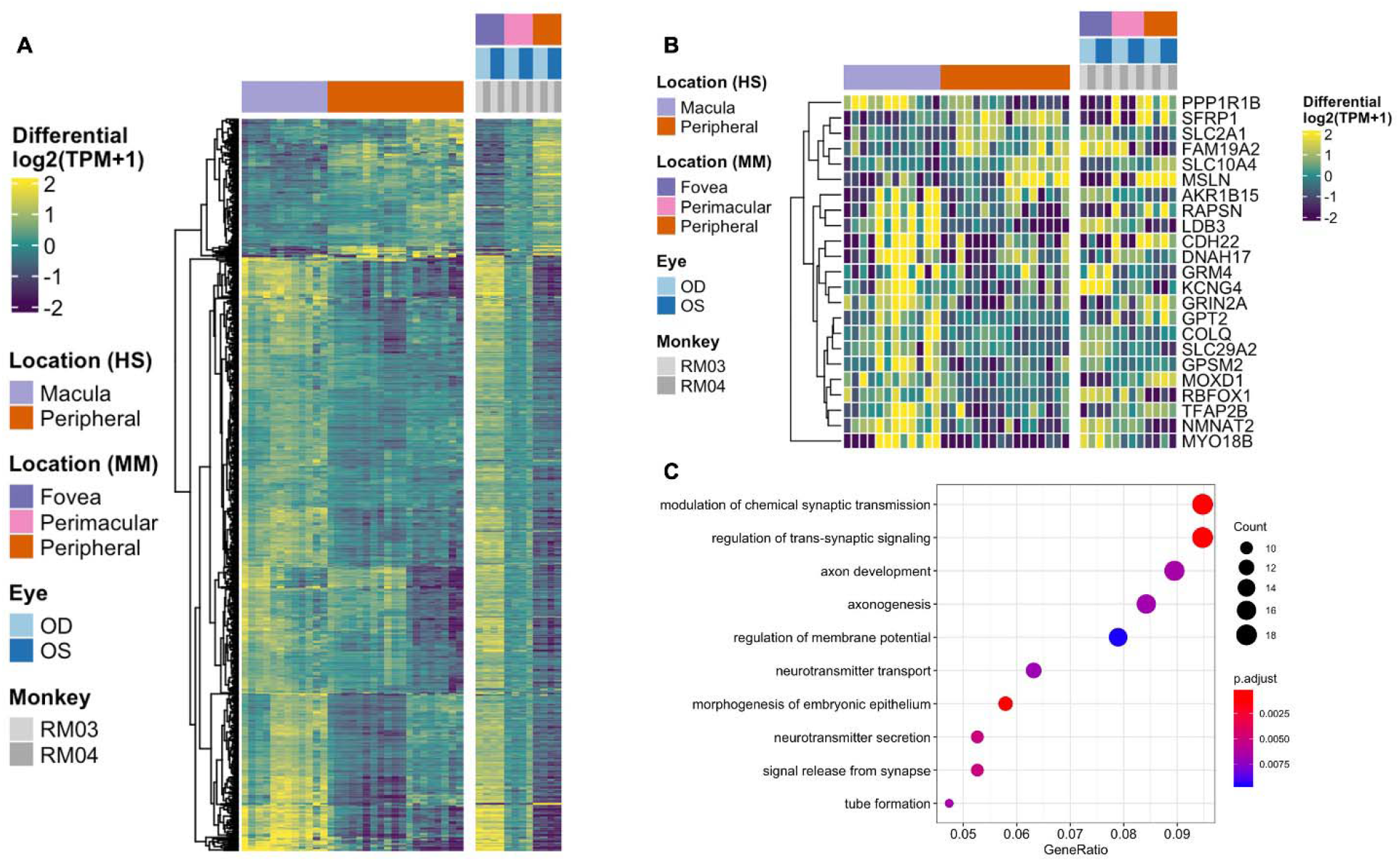
(A) Heatmap displaying the comparison of differential gene expression in the neural retina between published data from a human macula vs periphery set (Li et al., 2014, Whitmore et. al, 2014) and the macaque data presented here, with 664 genes following similar expression patterns. (B) A comparison of conserved differential expression to the human studies in the RPE/choroid tissue. Fewer genes follow similar patterns of differential expression. (C) GO enrichment for conserved differentially expressed genes between human and macaque datasets in the neural retina. Terms including neuronal connectivity and cellular morphogenesis are highly enriched.

To test whether there was any correspondence between our macaque tests and an independent single cell RPE cells taken from either the fovea or periphery, we extracted 219 cells from the scEiaD resource at plae.nei.nih.gov. We found ten genes that overlap between our macaque resource and the scEiaD resource, including *VIM*, *S100B*, and *PMEL*. Of these overlapping genes, *CXCL4*, *SULF4*, and *WFDC1* are shown to be enriched in the macula (Figure 4).

**Figure 4.**
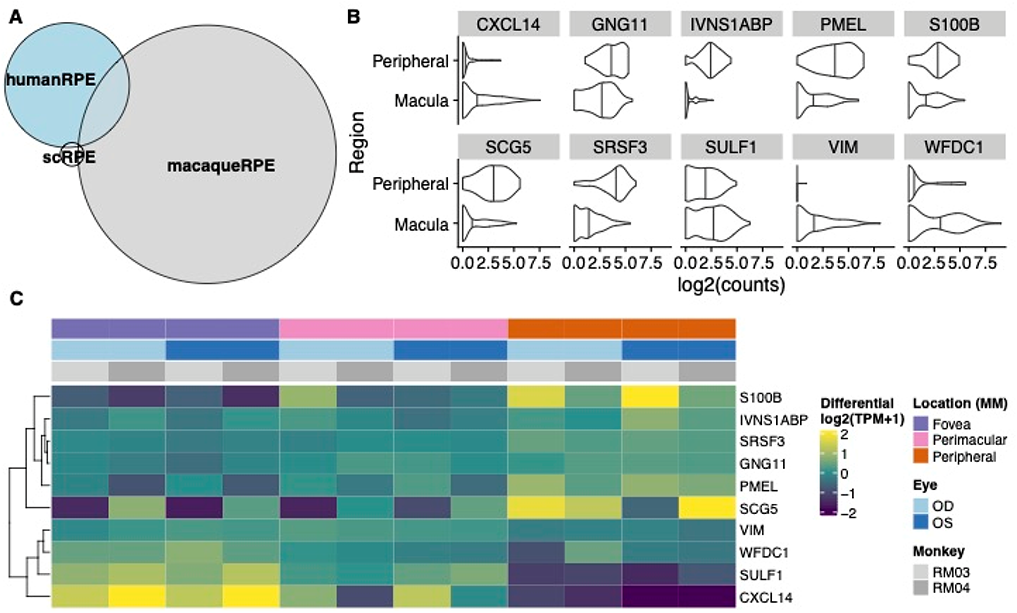
(A) Venn diagram of overlaps between differentially expressed genes between the fovea and periphery in three systems: human single cell RPE, bulk human RPE-choroid, and bulk macaque RPE-choroid. (B) Violin plot of the single cell expression with the ten genes which overlap between the human single cell RPE testing and bulk macaque RPE-choroid testing. (C) Heatmap of the same ten genes in our macaque RPE-choroid punches.

## Discussion

Using the Rhesus macaque as a model to study the transcriptome landscape of higher primate retina, we conducted RNA-sequencing across neural retina and RPE/choroid tissue layers. The small RNA punch size and bulk processing of the tissues allowed for high resolution and sensitivity of transcript quantification due to the decreased likelihood of variability from extended amplification, gene dropouts, and biological noise. Furthermore, it allows for the ability to profile rare and low-expressed transcripts as compared to single-cell RNAseq such as the 10x Genomics Chromium platform, in which only the top 20-30% of expressed genes are captured, or Drop-seq, which requires a more manual approach. Unlike previous studies, our approach included the perimacular retina to define contiguously changing gene expression across the retinal landscape, thereby reflecting the stepwise differences in the ratio of cone and rod composition. By doing this, we isolated the critical pathways regulating changes in retinal connectivity and RPE/choroid function necessary to support different photoreceptor mosaics.

The macaque gene expression changes observed in our data are orthogonally confirmed in human datasets. Furthermore, our findings also indicate conservation of contiguously changing gene expression patterns from the central to peripheral retina between Rhesus monkeys and humans, establishing a more available model system for future human transcriptomic studies. Rhesus monkeys offer an accessible model in which conditions of tissue acquisition can be further controlled, including age, optimization of biopsy and RNA extraction, and reduced time from enucleation to tissue dissection and RNA extraction.

PCA of gene expression data by tissue type predicts the distinct nature of the neural retina and RPE/choroid tissues, and further clustering by location confirms there are transcriptomic differences between regions of the retina. The grouping of transcriptomes by animal highlights the genetic variability observed between individuals. The lower number of differentially expressed genes in fovea vs perimacular comparisons versus fovea vs periphery indicates a continuum of gene expression. The highly variable patterns of gene expression across the neural retina corresponds to the changing composition of cell types based on location.

In contrast, the presence of distinct changing RPE/choroid gene expression suggests variability in the cell processes occurring across the retina in the RPE and choroid. Due to the relatively unchanged morphology of these tissues across the retina save for increased RPE diameter and choroidal thickness in the macula, the contiguous changes in gene expression observed may be a factor in driving changes in cell processes across the entire retina. GO terms enriched for the contiguously changing genes in the retina involved neuron projection and axon development as well as regulation of cell morphogenesis, supporting known findings that there are differences in circuitry and cone packing across the neural retina[10]. Furthermore, the changing gene expression across the RPE/choroid is enriched for GO terms such as cell junction organization and epithelial cell development, supporting reported findings that RPE diameter is smaller in the fovea/macula as compared to the periphery[38]. It is known that RPE cells form tight junctions, creating the outer blood-retinal barrier. These tight junctions serve as regulators of cell proliferation, polarity, and transport, as well as transducers of signals responsible for regulating cell size and shape[39]. Therefore, genes involved in cell junction organization and epithelial cell development may play a role in varied cell sizes across the RPE. GO terms involving urogenital and renal system development are also enriched in the RPE/choroid gene list, and interestingly, there are several heritable genetic conditions that affect both the eye and kidney, and there is also a link between chorioretinal thinning and chronic kidney disease, thought to be due to inflammation and endothelial dysfunction[40, 41]. In addition to observing greater levels of gene expression variability across location in the RPE-choroid than expected, several disease-relevant genes known to play roles in various ocular disorders including *TIMP3* (Various retinopathies[42]), *ABCA4* (Stargardt disease[12]), and *TYRP1* (oculocutaneous albinism[43]), were found to be significantly differentially expressed by location.

Differential expression of particular transcription factors, including neural retina- and RPE/choroid show conserved patterns of expression between macaque and human datasets, with *IRX2* being upregulated in the macula. Other differentially expressed transcription factors that show conservation with past human studies include *FOXI3*, which is upregulated in the periphery, and *POU4F2*, *POU4F1*, as well as its target, *RIT2*, all of which are enriched in the macula in both monkey and human datasets^7^. Additional transcription factors following similar gene expression patterns across species in the RPE/choroid include *VEZF1* and *NR2F1*, which are thought to be involved in retinal and vascular development[44, 45]. Identifying conserved location-specific transcription factors can provide insight into the regulatory landscape of the two tissues and how they relate to patterns of differential gene expression. It may be inferred then, that these changes in global gene expression across location and tissue, driven by distinct sets of transcription factors, likely reflect location-specific cellular processes and interactions.

Beyond patterns of transcriptional regulators, location-based tissue-tissue interactions are also indicative of specific pathways being up- or downregulated by location, suggesting specific roles in different locations of the tissues. The enrichment for Wnt signaling in kidney disease in the downregulated interactions in the macula may be connected to the GO enrichment for renal development in the RPE/choroid. Furthermore, ErbB signaling and peptide G-protein coupled receptor signaling are required for retinal development, with ErbB involved in neural crest development and adult pigment formation[46] and GCPR signaling involved in light processing in photoreceptors[47]. The tissue-tissue interaction analysis suggests these pathways are upregulated in the peripheral tissues of the eye. Certain pathways were enriched for tissue-tissue interactions both up- and downregulated in the macula, including Hippo signaling. Hippo-YAP signaling is known to play roles in both ocular development as well as disease, as it regulates retinogenesis, photoreceptor cell differentiation, and retinal vascular development, but misregulation also has associations with coloboma and optic fissure closure, uveal melanoma, and retinal degeneration[48]. The widespread enrichment of this pathway informs the many roles it plays in regulating the retinal landscape across location.

The three-way comparison of single cell, bulk human, and Rhesus macaque data showed many more differentially expressed genes detected in our macaque dataset than the other two sequencing methods. Furthermore, identifying differentially expressed genes in the Rhesus monkey data that intersect with the single cell data allows us to identify known RPE-specific genes that are conserved between the datasets including VIM[39], and PMEL[49], which we have shown to change contiguously by location. Of the three macula-enriched genes in the overlapping set, CXCL14 and WFDC1 have been previously identified as enriched in the macula compared to peripheral retina via RNA microarray[50], and WFDC1 enrichment in the macula was independently confirmed by both expression and immunostaining[51]. As such, our data shows the known macula-enrichment of WFDC1 to be conserved. As a serine protease inhibitor, mediator of endothelial cell migration and promoter of angiogenesis, its enrichment in the RPE and specifically, the macula, may suggest a role for WFDC1 in age-related macular degeneration [52], in which neovascularization beneath the macula is characteristic[50].

Considering the differential gene expression patterns across contiguous regions of the neural retina and especially the RPE/choroid allows for a combined approach in which we assessed the drivers of change as well as gross changes in gene expression by location. We identified highly enriched gene ontologies associated with each tissue location and layer, highlighted sets of contiguously changing transcription factors, determined important tissue-tissue interactions that highlight various up- and down-regulated location-specific pathways, and examined the conservation of gene expression patterns across multiple independent studies. In addition to the previous RNA-sequencing studies performed on human and non-human primate retinas, the data and findings presented here provide valuable resources for future studies aimed at identifying regional specialization of the retina and understanding disease mechanisms.

## Acknowledgements

We would like to thank Bob Wurtz, Bruce Cumming, James Cavanaugh, Mitch Smith of the Laboratory of Sensorimotor Research, National Eye Institute for providing non-human primate samples. We would also like to thank the NEI/NIMH Animal Program, including Jim Raber and Ginger Tansey, the National Institutes of Health Intramural Sequencing Center, where RNA sequencing was performed, and Dr. Sheldon Miller for helpful comments and advice about the experiments and manuscript.

## Supplemental Data

**Figure S1.**
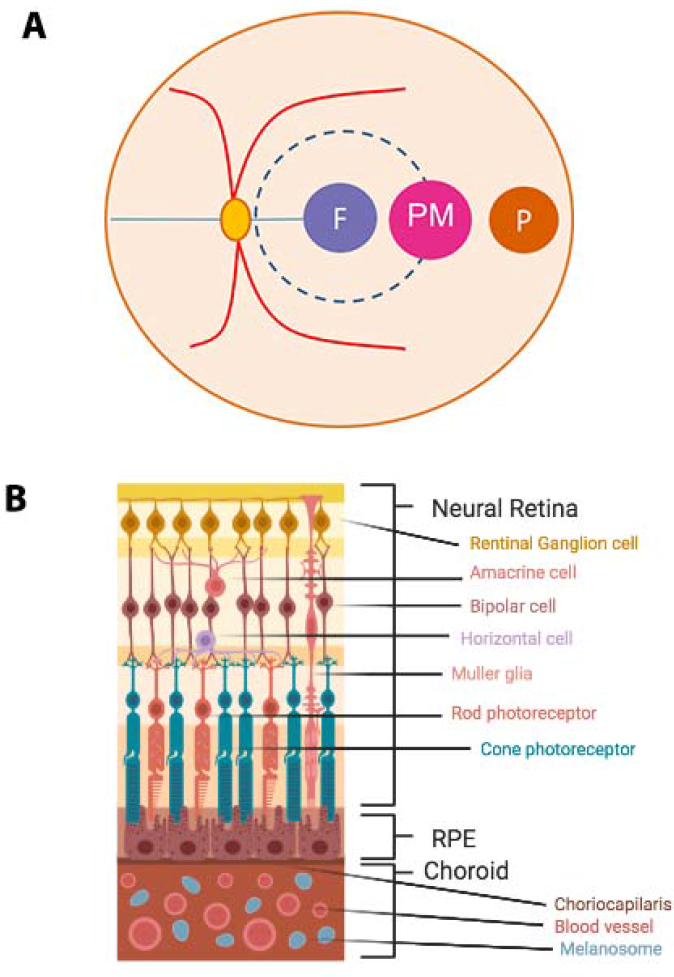
(A) Layout of punch biopsies taken from the Rhesus monkey eyes, including foveal (F), perimacular (PM), and peripheral (P) samples. (B) Diagram of the retina, highlighting the cell types in the neural retina as well as the RPE and choroid tissues (made in Biorender).

**Figure S2.**
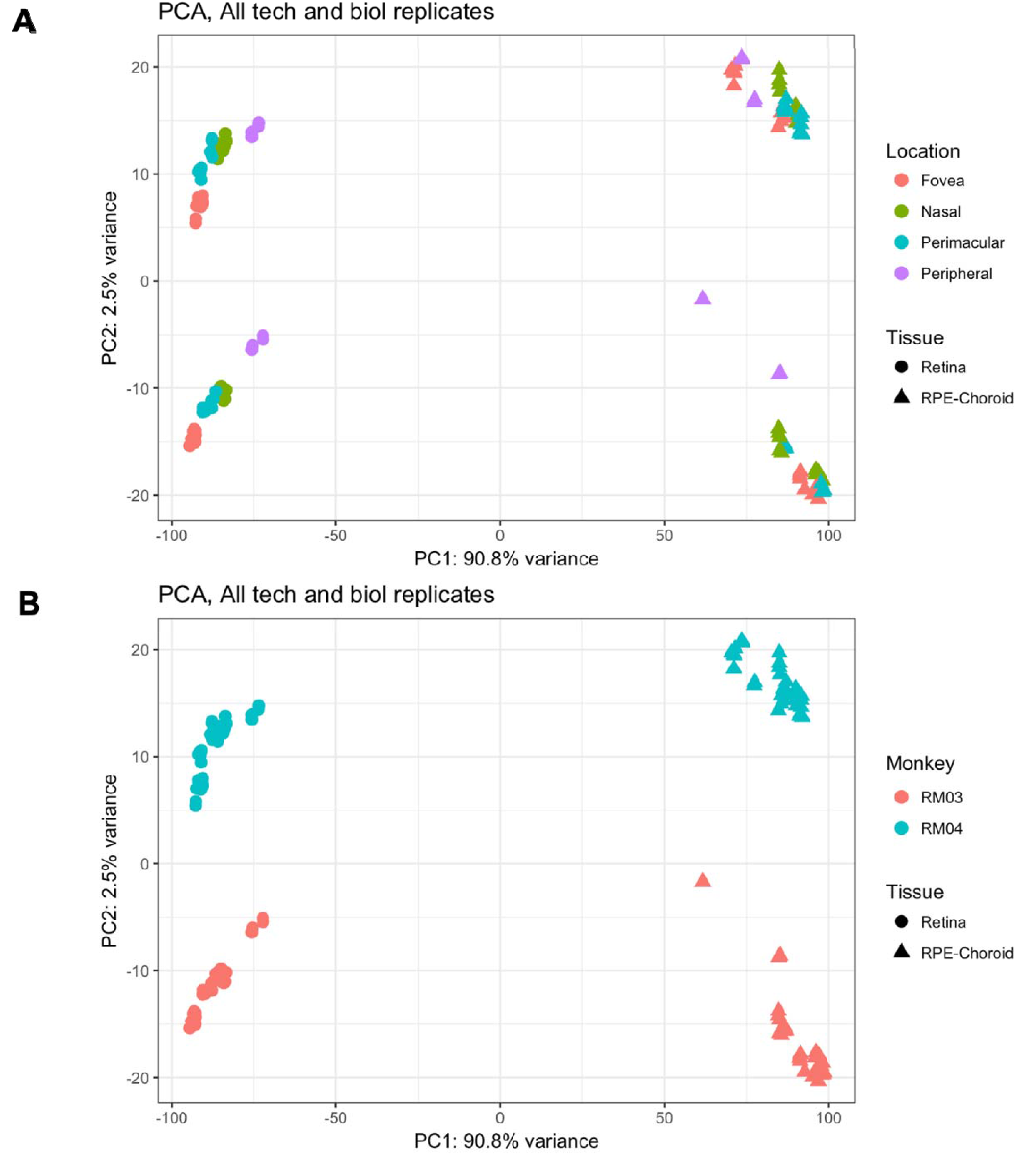
PCA clustering for the 1000 most variable genes sequenced. (A) Samples primarily cluster by tissue type, with RPE/choroid samples clustering away from retina samples. (B) Within the retina and RPE/choroid, foveal and peripheral samples tend to cluster furthest apart with perimacular samples clustering less tightly between.

**Figure S3.**
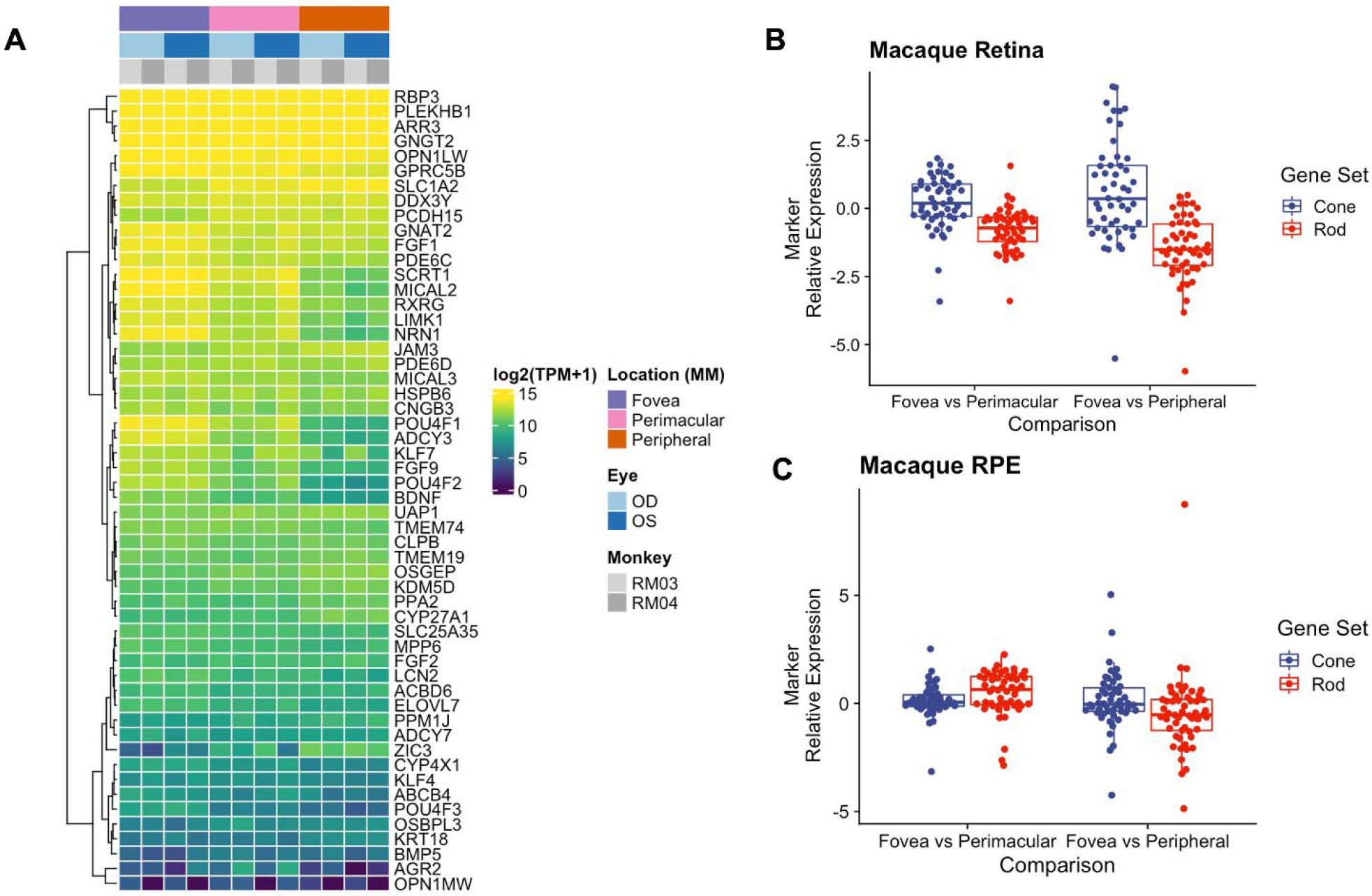
(A) Heatmap depicting macaque expression of published rod-enriched gene expression (Holt et. al 2015, Mustafi et. al, 2016) across the neural retina reflects cellular composition of rods in the retina. Genes are highly expressed in the periphery, where rod composition is highly enriched, and lowly expressed in the fovea/macular which is highly cone-enriched. (B) Peripheral and perimacular retinal samples express rod-enriched genes at a 2.8- and 1.7-fold increase compared to foveal retinal samples. The periphery is 1.6-fold enriched in cone signal compared to the fovea in the neural retina (Table S1). (C) Changes in rod- and cone-enriched gene expression do not vary greatly by location in RPE/choroid.

**Figure S4.**
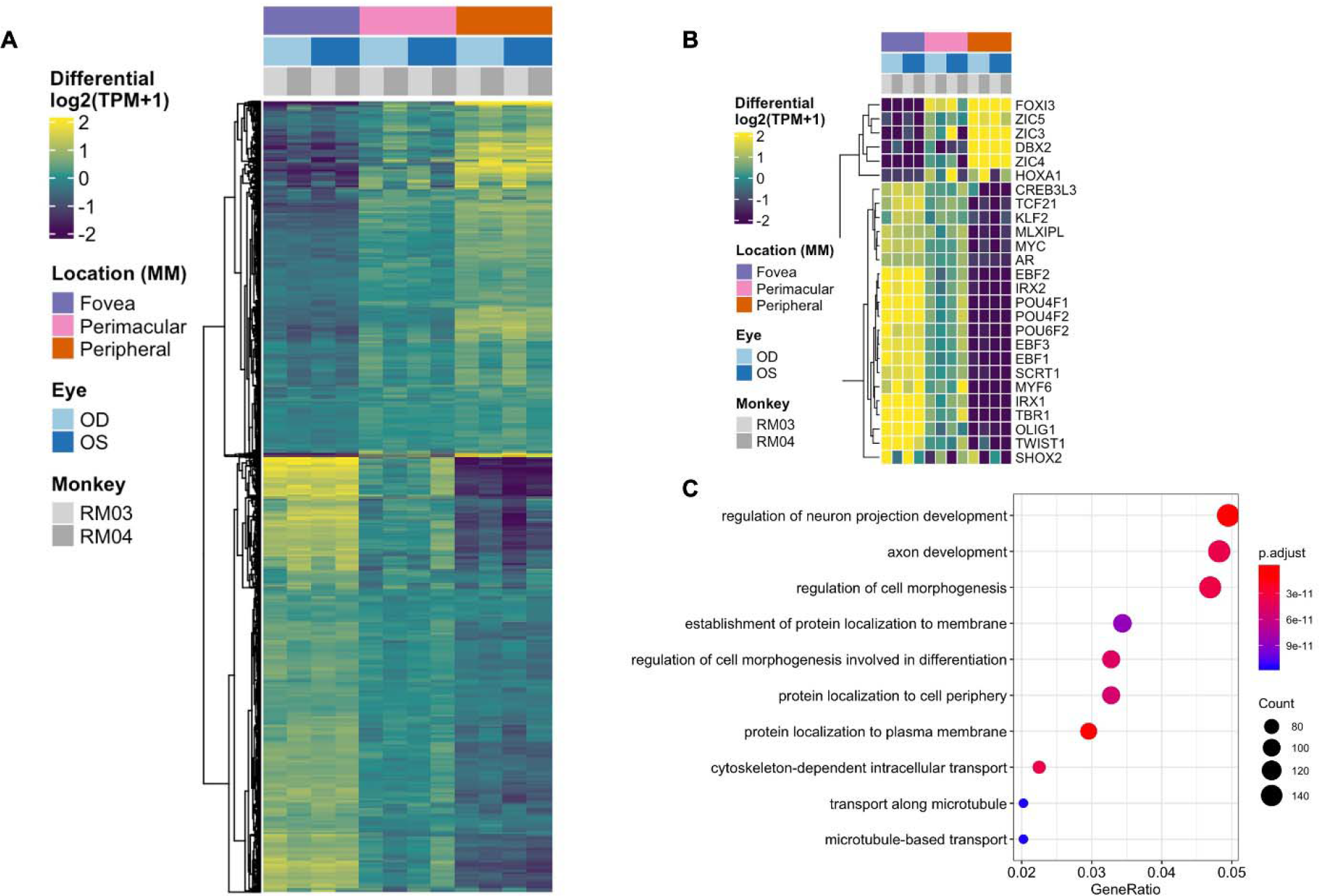
(A) Heatmap showing contiguous differential expression across location of the neural retina tissue, also exhibiting distinct blocks of progressive up and down regulated gene expression as seen for RPE/choroid in Fig. 1C. (B) A subset of genes identified as differentially expressed transcription factors by location in the neural retina. Changes in transcriptional regulators can be a factor in driving gene expression changes. *These transcription factors have published associations with the respective tissue in the literature. (C) Top 10 enriched gene ontology (GO) terms from the retina differentially expressed gene set, including neuron projection, axon development, regulation of cell morphogenesis, and protein localization and transport.

**Figure S5.**
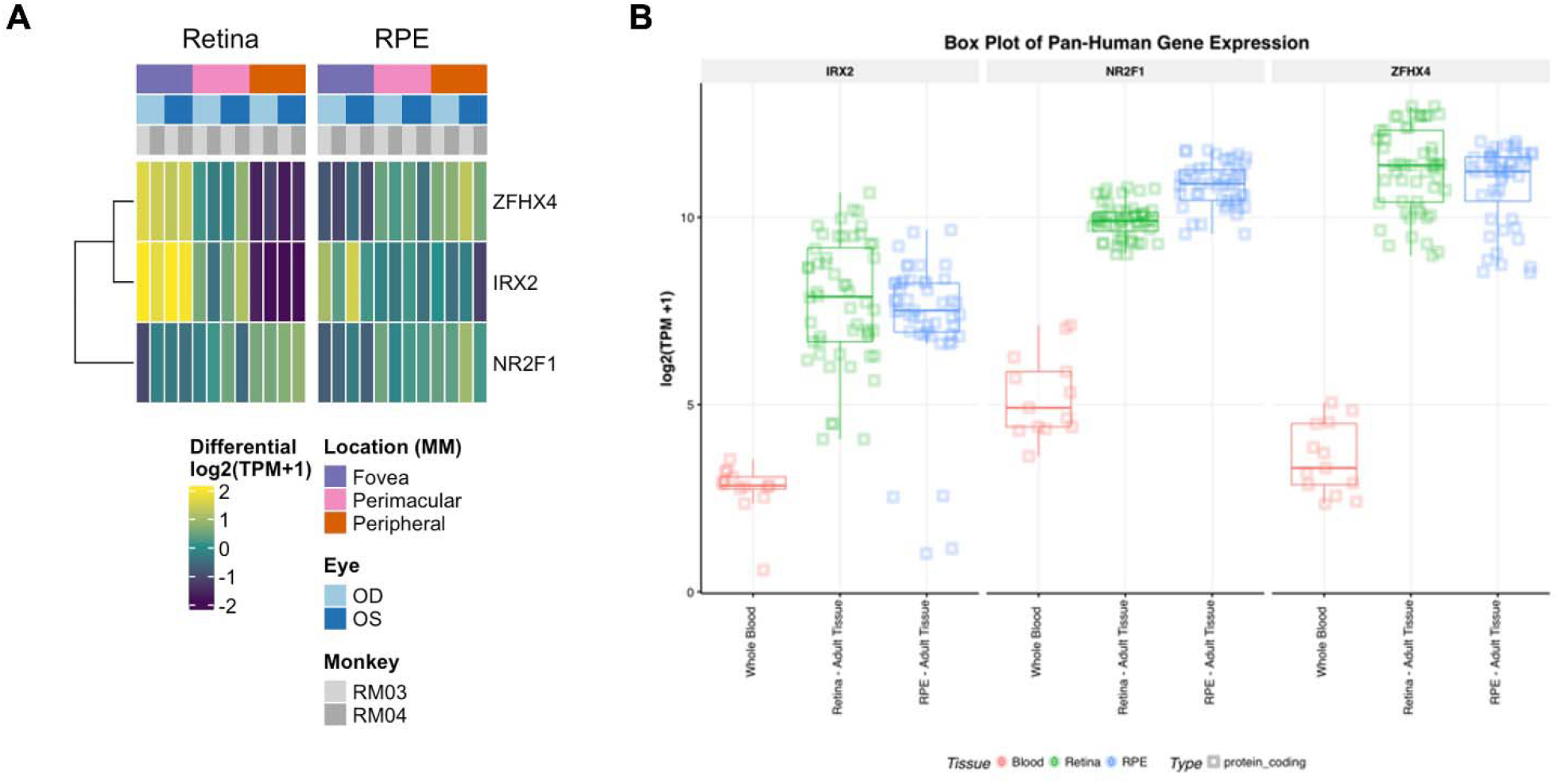
(A) Expression of commonly expressed transcription factors between the neural retina and the RPE/choroid. While *NR2F1* and *IRX2* follow similar expression patterns across location, *ZFHX4* actually displays opposite expression patterns in the retina and RPE/choroid tissues. (B) Expression of these three transcription factors from the publicly available EyeIntegration database (Swami and McGaughey et al., 2019) showing similar expression levels of each gene in both tissue types with whole blood for comparison.

**Figure S6.**
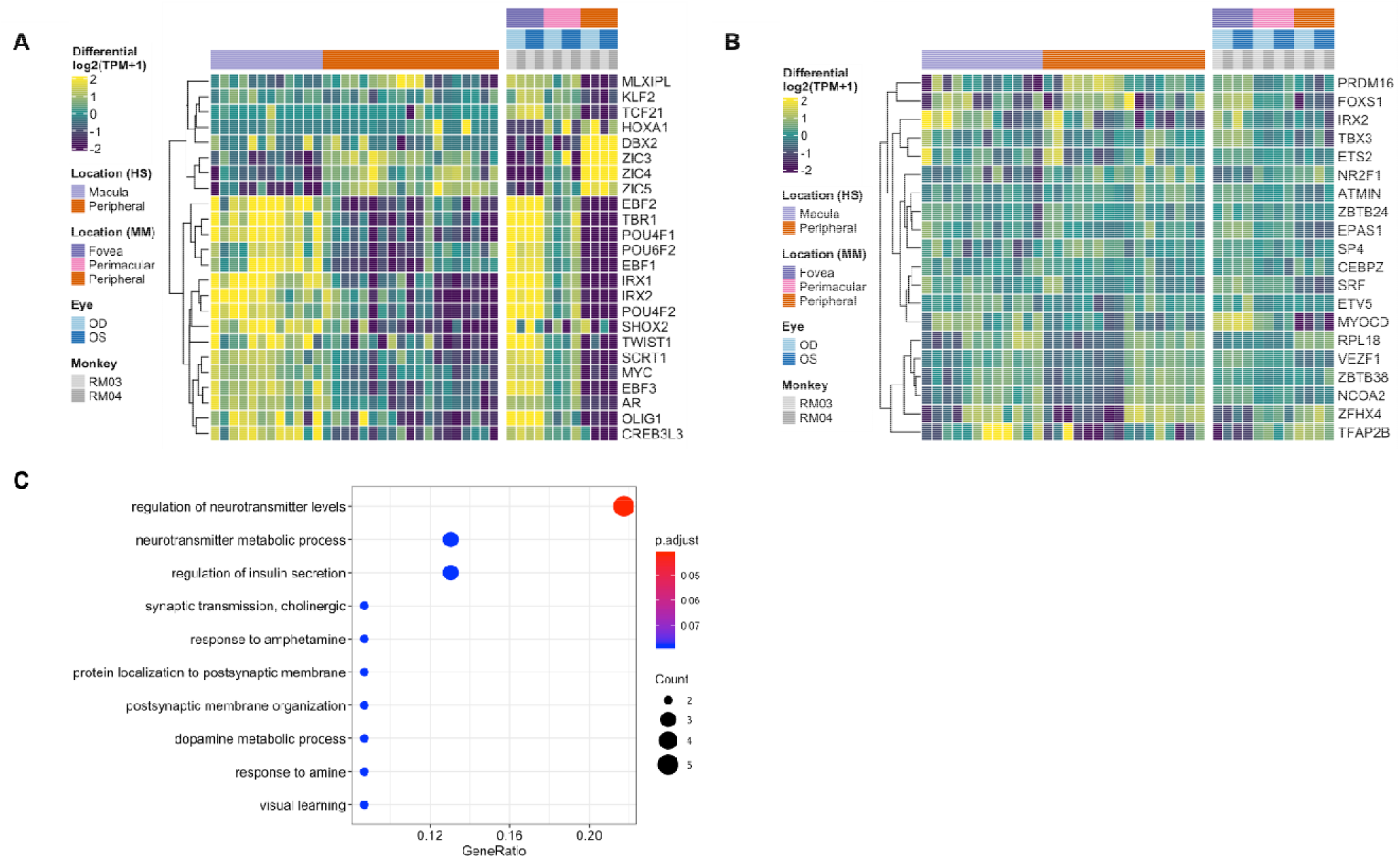
(A) A comparison of conserved transcription factor differential expression in the neural retina between published data from a human macula vs periphery set (Li et al., 2014, Whitmore et. al, 2014) and the macaque data presented here. (B) A similar comparison of human conservation of transcription factor differential expression in the RPE/choroid tissue. (C) Top enriched GO terms for the conserved gene expression across fovea to periphery in the RPE/choroid. Top terms include regulation of neurotransmitter levels and metabolic processes as well insulin secretion.

**Figure S7.**
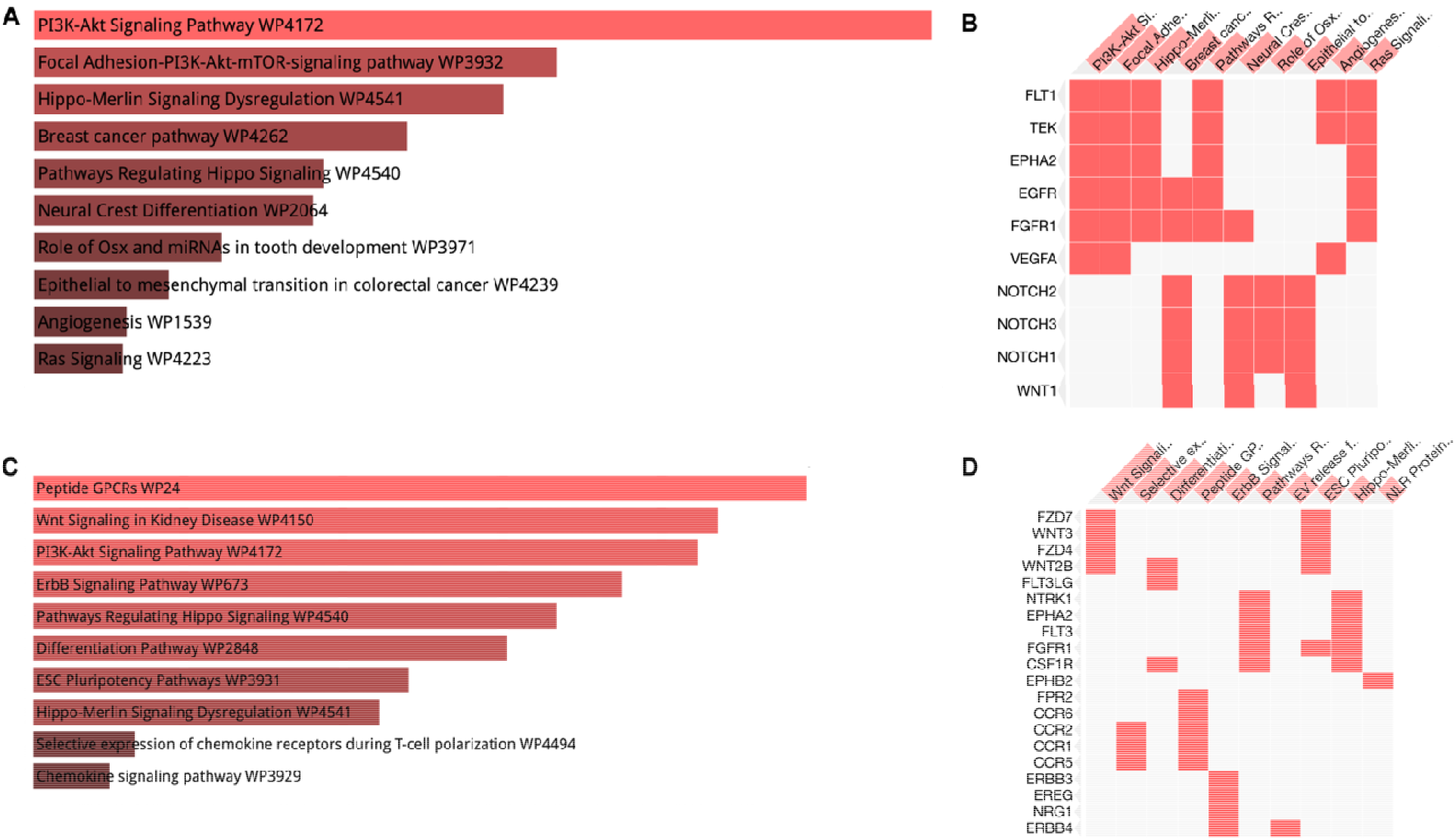
(A) Enrichr bar graph depicting highly enriched pathways via the WikiPathways 2019 Human database for the downregulated ligand/receptor interactions in the macula. Top enriched pathways include Wnt signaling in kidney diseaseand ErbB signaling. (B) Enrichr clustergram depicts the genes contributing to the enriched pathway terms including *WNT*, *FZD4*, and *ERBB3*. (C) Enrichr bar graph for upregulated ligand/receptor interactions in the macula. Top enriched pathways include PI3K-Akt signaling and Focal Adhesion-PI3K-Akt-mTOR, both of which are involved in the VEGF signaling pathway. (D) Enrichr clustergram is enriched for genes including *VEGFA*, *FLT1*, and *KDR*, all of which are key components of VEGF signaling.

**Table S1.** Rod- and Cone-Enriched gene expression in Retina and RPE/choroid compared across location

| Tissue | Comparison | GeneSet | estimate | enrichment | statistic | p.value | parameter | conf.low | conf.high |
| --- | --- | --- | --- | --- | --- | --- | --- | --- | --- |
| Retina | Fovea vs Perimacular | Rod | -0.7821306 | -1.7196686 | -7.4071987 | 1.11E-09 | 52 | -0.9940137 | -0.5702476 |
| Retina | Fovea vs Peripheral | Rod | -1.4620671 | -2.7550282 | -8.9989078 | 3.48E-12 | 52 | -1.7880902 | -1.1360439 |
| Retina | Fovea vs Perimacular | Cone | 0.20531647 | 1.15293923 | 1.46077663 | 0.15045974 | 49 | -0.0771353 | 0.48776821 |
| Retina | Fovea vs Peripheral | Cone | 0.6492966 | 1.56840332 | 2.441338 | 0.01829112 | 49 | 0.11483133 | 1.18376187 |
| RPE | Fovea vs Perimacular | Rod | 0.47748612 | 1.39231546 | 3.3946687 | 0.00132244 | 52 | 0.1952359 | 0.75973635 |
| RPE | Fovea vs Peripheral | Rod | -0.4613632 | -1.3768422 | -1.8370341 | 0.07192272 | 52 | -0.9653239 | 0.0425975 |
| RPE | Fovea vs Perimacular | Cone | 0.13785476 | 1.10026784 | 1.27906465 | 0.20689857 | 49 | -0.0787328 | 0.35444235 |
| RPE | Fovea vs Peripheral | Cone | 0.14454836 | 1.10538455 | 0.76617502 | 0.44724757 | 49 | -0.2345828 | 0.52367948 |

**Table S2.**
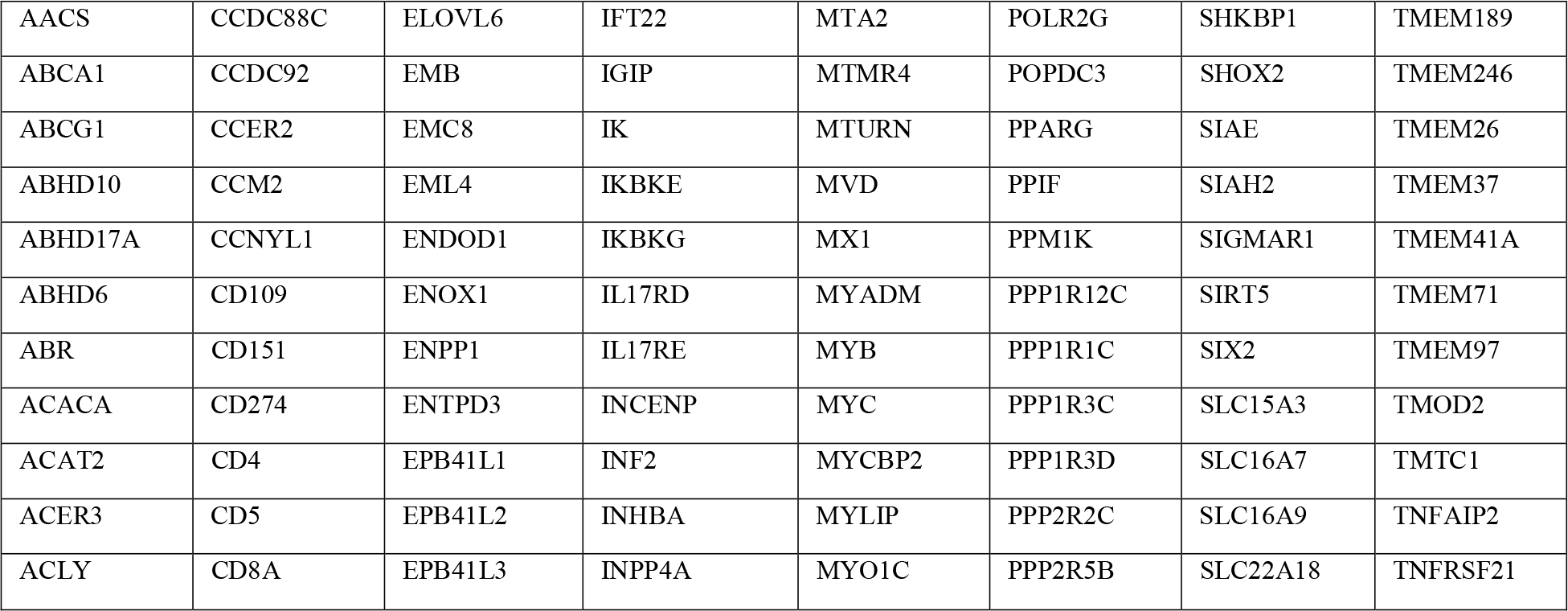
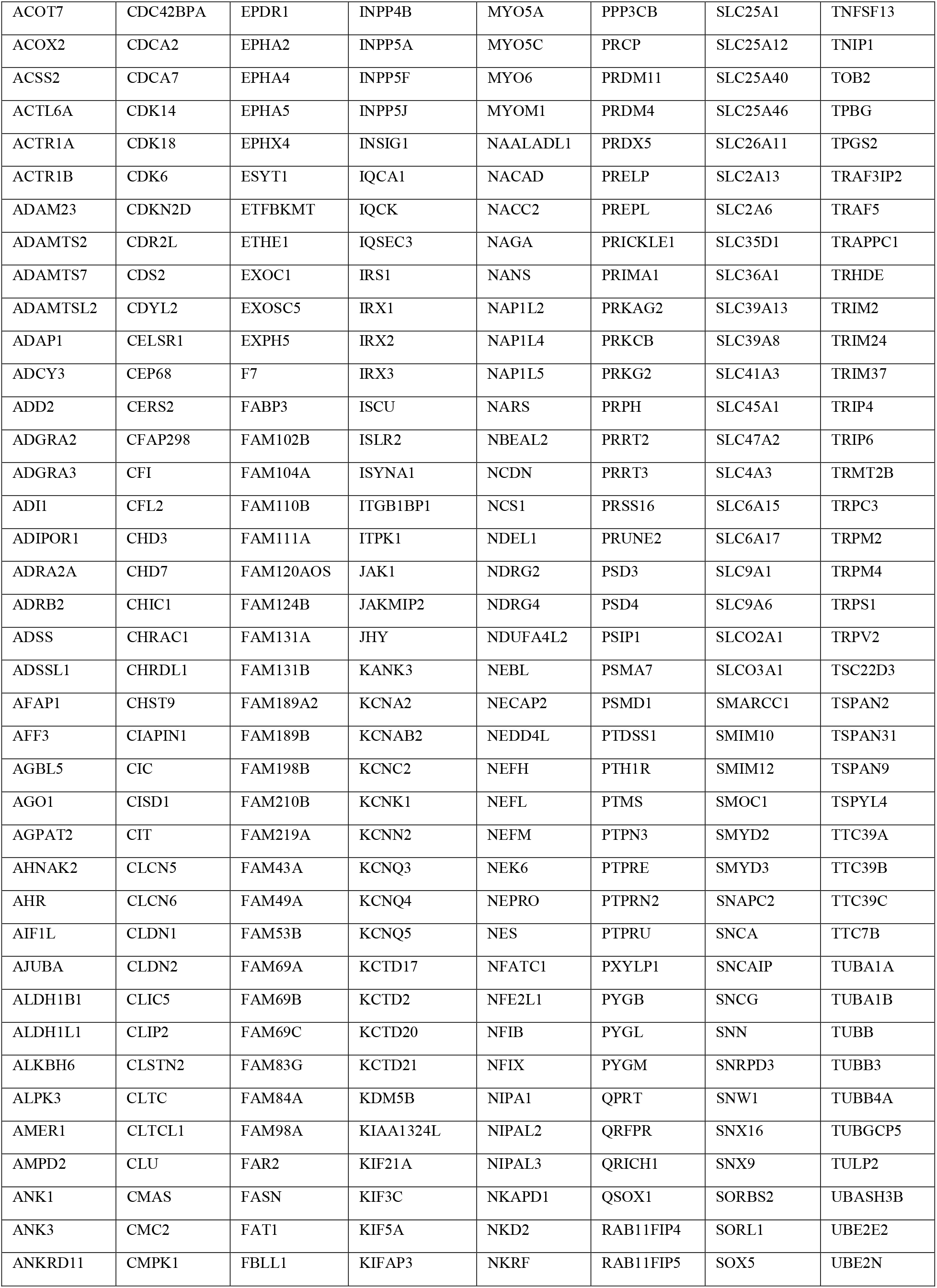
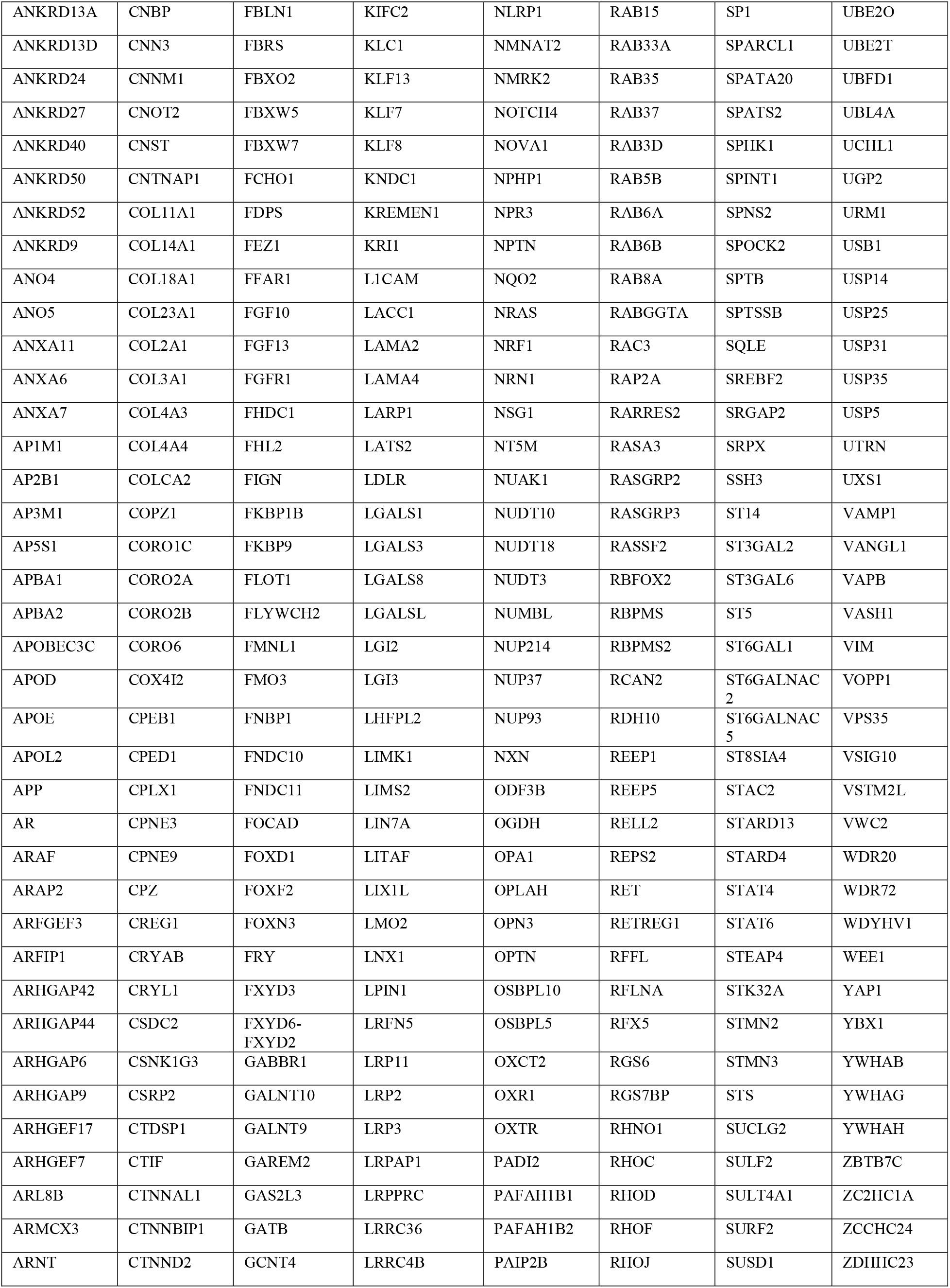
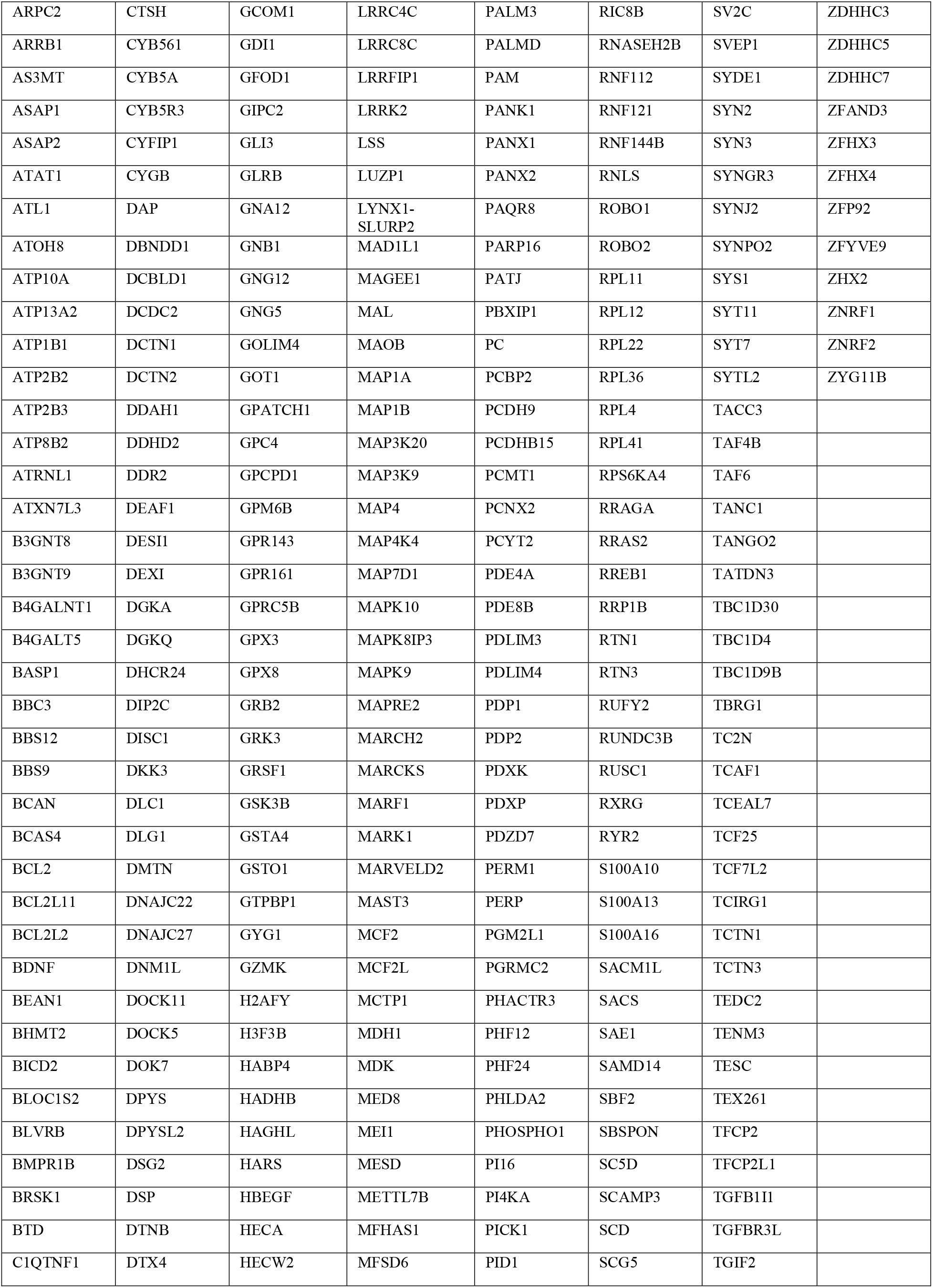
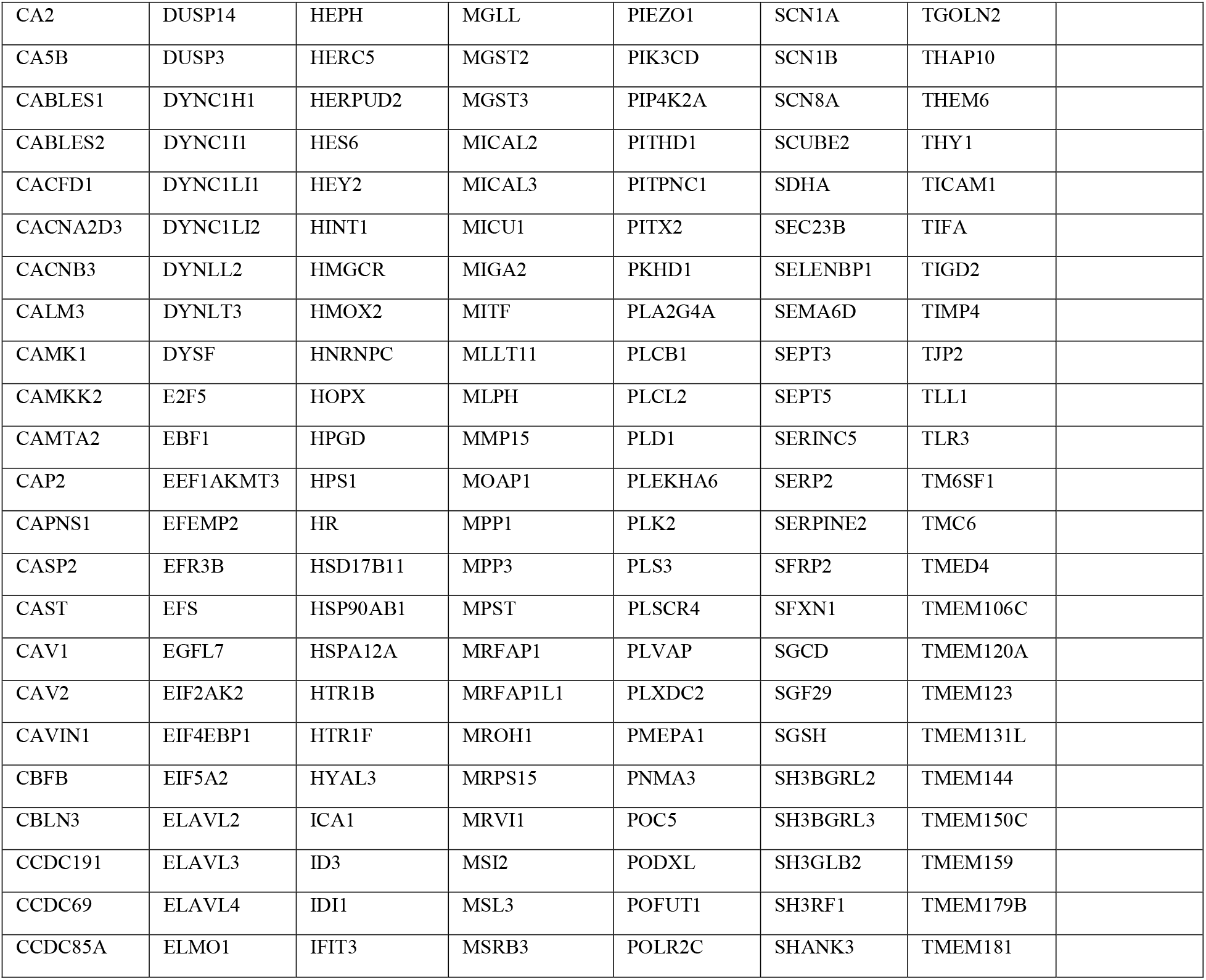
List of conserved genes between human and macaque data as seen in the heatmap (Fig. 3A)

